# Impairment of neuronal mitophagy by mutant huntingtin

**DOI:** 10.1101/330001

**Authors:** Sandra Franco-Iborra, Ainhoa Plaza-Zabala, David Sebastian, Miquel Vila, Marta Martinez-Vicente

## Abstract

Neuronal homeostasis depends on the proper functioning of different quality control systems. The accurate degradation of dysfunctional mitochondria by mitophagy is essential for maintaining mitochondrial quality and quantity in neurons. Huntingting protein (Htt) can interact with numerous other proteins and thereby perform multiple biological functions in the cell. We investigated the role of Htt during neuronal mitophagy in post-mitotic differentiated striatal neurons avoiding artificial overexpression of mitophagy-related proteins. Our study shows that Htt promotes the physical proximity of different protein complexes during initiation of the mitophagy process. Htt is needed to promote the ULK1 complex activation, the recruitment of the essential receptors of mitophagy –optineurin and NDP52- and the interaction of these receptors with LC3-II. Mutant Htt also causes a gain of toxic function avoiding the formation of the Beclin1-Vps34 initiation complex. We report that the presence of the expanded polyQ tract can affect the efficiency of neuronal mitophagy and consequently promote the accumulation of damaged mitochondria and increase oxidative stress, leading to mitochondrial dysfunction and contributing to neurodegeneration in Huntington’s disease.

## INTRODUCTION

Huntington’s disease is an autosomal-dominant neurodegenerative disorder caused by an expansion of the CAG trinucleotide repeat encoding a polyglutamine (polyQ) tract in the amino-terminal region of Huntingtin (Htt) protein [1]. HD is characterized by motor dysfunction, cognitive decline and psychiatric disturbances. Despite the ubiquitous expression of mutant Htt (mHtt), this disorder is characterized by a preferential atrophy of the striatum caused by the loss of GABAergic medium spiny neurons in the caudate nucleus and putamen, although other brain areas such as the cerebral cortex are also affected [2,3]. The neuropathological hallmark of HD is the formation of inclusion bodies within the affected neurons due to the presence of an expanded polyQ tract that renders mHtt prone to aggregation [4]. Htt is a highly conserved and essential for mammalian embryonic development [5–7]. Adult loss of Htt leads to neurodegeneration, demonstrating Htt’s essential role in maintaining neuronal function and survival [8]. Htt expression is ubiquitous and the protein is present in many subcellular compartments [9–11]. Htt is a particularly large protein (347.603 kDa) with multiple protein-protein interaction regions, consequently several interaction partners have been described [10–12], suggesting that Htt acts as a scaffold to stabilize different protein complexes. To date, Htt has been proposed to be involved in transcriptional regulation [13], axonal trafficking of vesicles, autophagosomes and mitochondria [14–16], nuclear export/import, ubiquitin-mediated proteolysis and regulation of the endosomal and autophagic systems among many other cellular mechanisms [17].

Cellular homeostasis depends on the proper functioning of different quality control systems that ensure the continuous turnover, degradation and recycling of all intracellular components. Autophagy is the catabolic mechanism by which intracellular components such as proteins, organelles and aggregates are delivered to lysosomes for degradation [18,19]. This mechanism plays an essential role in neuronal homeostasis and survival since neurons, in contrast to other non-postmitotic cells, require a constitutive basal autophagic activity to prevent the accumulation of misfolded and aggregated proteins, eliminate unwanted organelles and keep the cell ‘clean’ [20,21].

Dysfunction of the cellular proteolytic systems has been extensively linked to HD. In addition to impairment of the proteasome system [22], different forms of autophagy are also affected in HD. Defective macroautophagy has been reported in HD-affected neurons [23,24] and linked to the basal upregulation of chaperone-mediated autophagy (CMA) as compensatory mechanism of both ubiquitin-proteasome system (UPS) and macroautophagy failure [25].

Mitophagy is a form of selective macroautophagy in which damaged mitochondria are targeted for degradation in the lysosome. While Basal mitophagy is responsible for the constant turnover of the mitochondrial pool, mitophagy can also be induced during development in certain cell types or as a stress-response mechanism following mitochondrial membrane depolarization or hypoxia [26]. Despite the different molecules involved, the various forms of mitophagy follow a similar mechanism where an adapter or receptor molecule acts as bridge between the targeted mitochondria to be eliminated and the LC3-II protein present in the autophagosome membrane. This permits the targeted mitochondria to be engulfed by the autophagosome and subsequent degradaded after fusion with a lysosome. The ability of Htt to physically interact with several partners provides structural stabilization to different protein complexes and allows physically proximity required to carry out different cellular mechanisms. Recently it was shown that Htt can act as a scaffold protein during the initiation of stress-induced selective autophagy [27,28]. To date Htt has been shown to interact with unc-51 like kinase 1 (ULK1) and p62/SQTM12. ULK1 is required to initiate macroautophagy while p62 is one of the selective macroautophagy receptors that binds simultaneously to autophagosomes via the interaction with LC3-II and a substrate through the direct interaction with polyUB chains tagged to the substrate [27,28].

Here, we investigated the genuine role of Htt in neuronal mitophagy, not only through its interaction with ULK1 and p62, but also through its interaction with other essential components of mitophagy mechanisms. In contrast to previous works, we used differentiated striatal neurons to study Htt’s role in neuronal mitophagy without artificially modifying the endogenous levels of the molecular components involved in this mechanism. Furthermore, to promote stress-induced mitophagy, we achieved severe mitochondrial depolarization with the mitochondrial uncoupler CCCP as frequently used in previous reports. We also studied mitophagy induction in response to mild depolarization with low concentrations of rotenone. This non-massive mitochondrial depolarization more closely mimics the typically moderate mitochondrial damage that occurs during neurodegeneration.

## RESULTS AND DISCUSSION

### Mitophagy is affected in ST-Q111 cells

Wild type (WT) Htt has been implicated at the cargo recognition step in stress-induced selective autophagy due to its ability to act as a scaffold that brings together different proteins required for autophagy to take place [27,28,23]. To study the consequences of glutamine expansion in huntingtin’s role in neuronal mitophagy, we using differentiated striatal neurons derived from a knock-in mouse expressing human exon 1 carrying a polyQ tract with 7 glutamines STHdh-Q7 (ST-Q7, control) or 111 glutamines STHdh-Q111 (ST-Q111, mutant) inserted into the endogenous mouse *Htt* gene. To begin with, we directly studied mitochondrial clearance inside the lysosomes of these neurons. Mitophagy can be monitorized by the colocalization of mitochondria within lysosomes. Under basal conditions, the number of lysosomes loaded with mitochondrial material in the presence of lysosomal inhibitors was similar between control and mutant striatal cells (**Fig. 1A**). However, after mild (rotenone 1 μM for 4h) and acute (CCCP 10 μM during 24h) mitochondrial depolarization, control cells showed a clear increase in the number of lysosomes colocalizing with mitochondria, but no increase was observed in the mutant ST-Q111 cells (**Fig. 1A**). The colocalization assays of the latter showed that a lower number of lysosomes were loaded with mitochondria, indicating that upon depolarization mHtt cells seem to be less-efficient in digesting damaged michondria. To further analyze mitochondrial clearance in our differentiated striatal cell line, we took advantage of the properties of Keima, a fluorescent protein with a pH-dependent excitation and which is relatively resistant to lysosomal degradation [29,30]. Keima can be expressed in mitochondria, where it exhibits a peak excitation at 440 nm. Upon delivery of mitochondria to lysosomes for degradation, mitochondrial matrix-localizing Keima (mitoKeima) undergoes a shift towards a 586 nm excitation wavelength due to the acidic lysosomal pH [29]. We transfected mitoKeima into differentiated ST-Q7 and ST-Q111 cells and induced a mild or acute mitochondrial depolarization. Under these conditions, mitochondria are delivered into lysosomes where Keima is present in the lysosomal lumen. The mitoKeima intensity upon red-laser excitation was then analyzed as a reporter of active mitophagy. An increase in mitoKeima intensity was observed in ST-Q7 cells, concomitant with the severity of the mitochondrial insult, indicating that mitochondria were being loaded into lysosomes upon depolarization. No changes were observed in ST-Q111 cells, where a tendency towards a decrease in mitoKeima intensity was actually observed, suggesting an alteration in the mitophagy process (**Fig. 1B**). As another measure of mitochondrial degradation, we quantified mtDNA as a reporter of mitochondrial mass [30]. To this end, we immunostained differentiated ST-Q7 and ST-Q111 cells with an anti-DNA antibody that preferentially stains mtDNA [30,31]. In ST-Q7 cells there was a loss of DNA intensity upon rotenone and CCCP treatment, indicative of mitochondrial removal in response to damage. However, no significant changes were observed in ST-Q111 cells, further implying that mitochondria were not being degraded with the same efficiency as ST-Q7 cells upon stress-induction (**Fig. 1C**). These results indicate that induced mitophagy is impaired in differentiated ST-Q111 cells, suggesting that mHtt might directly or indirectly affect the efficiency of this process. Indeed, we have previously described the presence of “empty” autophagosomes in HD striatals cells as a consequence of inefficient mitochondrial loading during selective macroautophagy that includes fewer engulfed mitochondria [23]. Other authors have also described reduced levels of mitophagy in a HD mouse model expressing mt-keima in vivo, confirming that this mechanism is altered in the presence of mHtt. Other authors have also described reduced levels of mitophagy in a HD mouse model expressing mt-keima in vivo, confirming that this mechanism is altered in the presence of mHtt [32]

**Figure 1.**
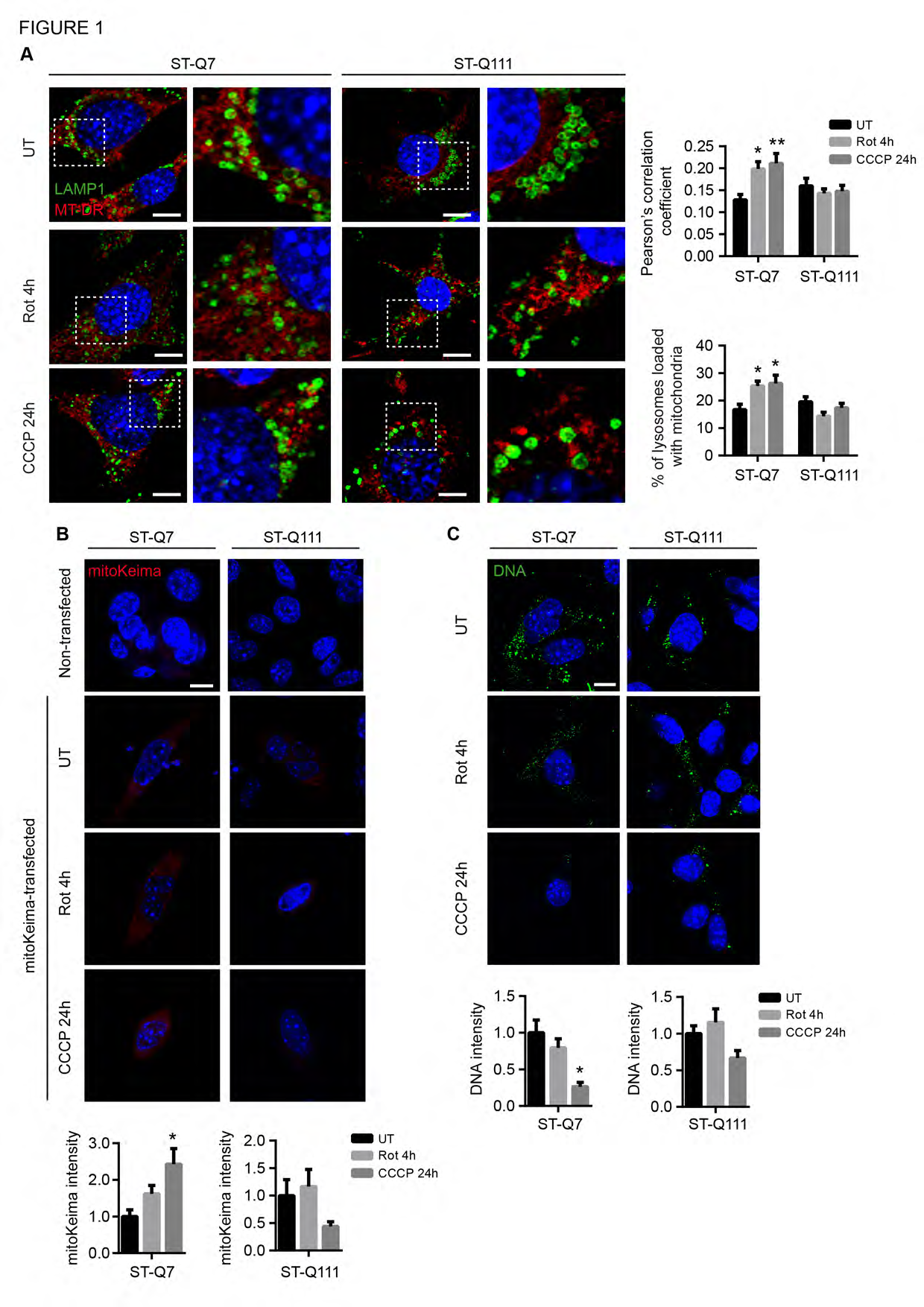
Mitophagy is affected in ST-Q111 cells. (**A**) Representative images of LAMP1 and MitoTracker^®^ DeepRed immunostained ST-Q7 and ST-Q111 cells untreated (UT) or treated with rotenone (Rot, 1 μM, 4h) or CCCP (10 μM, 24h). Insets show higher magnification images. Colocalization degree was quantified using the Pearson’s correlation coefficient and the ratio of LAMP1-stained lysosomes loaded with MitoTracker^®^ DeepRed-stained mitochondria. A minimum of 30 cells were analyzed per condition. (**B**) Representative images of non-transfected or mitoKeima-transfected ST-Q7 and ST-Q111 cells UT or treated with Rot (1 μM, 4h) or CCCP (10 μM, 24h). Lysosomal-positive mitoKeima signal was calculated as the ratio of intensity (594 nm-light excitation) per cell area, represented as the fold change compared with mitoKeima-transfected UT conditions. A minimum of 20 cells were analyzed per condition. (**C**) Representative images of ST-Q7 and ST-Q111 cells immunostained with anti-DNA UT or treated with Rot (1 μM, 4h) or CCCP (10 μM, 24h). Quantification is represented as the fold change in DNA intensity compared with UT conditions. 15 to 50 cells were analyzed per condition. (**A-C**) Scale bar: 10 μm. Data are presented as mean ± s.e.m. ^*^ P < 0.05, ^**^ P < 0.01 compared to UT condition after (**A**) two-way ANOVA followed by Tukey’s post hoc test (**B-C**) one-way ANOVA test followed by Tukey’s post hoc test.

### Mitophagy in differentiated striatal cells is not mediated by Parkin translocation to mitochondria

To gain insight into the mechanism by which mHtt affects mitophagy, we analyzed the different steps of the mitophagy process. According to many studies, the initiation of mitophagy commences with the retention of full-length PINK1 protein on the mitochondrial surface and the recruitment and activation of parkin protein [33–39]. Active parkin ubiquitinates a variety of substrates present on the outer mitochondrial membrane (OMM),either elongating pre-existing ubiquitin chains or ubiquitinating substrates *de novo*. Parkin-mediated ubiquitination feeds a positive feedback loop, amplifying the signals that lead to mitochondrial degradation [30]. We investigated whether PINK1 and parkin levels could be affected in ST-Q111 cells upon induction of mitophagy. We observed that PINK1 protein levels remained steady in differentiated ST-Q7 cells upon mild or acute depolarization; however, in differentiated ST-Q111 cells, basal PINK1 protein levels were slightly increased but again, no changes were observed upon rotenone- or CCCP-induced mitophagy (**Fig. 2A**). Depolarization induced by various effectors did not alter parkin protein levels in total cellular homogenates, but more importantly, we were unable to detect parkin protein translocation to mitochondria using two distinct methods. Parkin protein was not detected by immunoblot in isolated mitochondrial fractions; instead, all parkin protein remained cytosolic (**Fig. 2B**). Parkin translocation to mitochondria was also not detected by confocal immunofluorescence upon basal or depolarization-induced conditions in differentiated ST-Q7 and ST-Q111 cell lines (**Fig. 2C**). This scenario is similar to that observed in other neuronal cells where endogenous parkin translocation to the mitochondria was not detectable [40–43]. Indeed, most previous studies PINK1/parkin-dependent mitophagy have been performed with overexpression of tagged parkin protein since levels of endogeneous parkin were not detectable. To test if the lack of mitochondrial-associated parkin after depolarization was due to the lower levels of parkin protein, we artificially overexpressed parkin protein tagged to a fluorescent protein (EGFP). We could not promote a clear translocation of EGFP-parkin within a short time with mild depolarization (1 μM Rot 4h). Only after 24h with a strong depolarizing agent (10 μM CCCP) it was possible to observe a partial translocation of EGFP-parkin from a diffuse cytosolic distribution to a mitochondrial localization (**Fig 2D**). Nevertheless, EGFP-parkin mitochondrial translocation was only detectable in less than 50% of both ST-Q7 (42,7% ± 11,8) and ST-Q111 (46,3% ± 11,2) differentiated striatal cells (**Fig 2D**), as previously reported [44]. This contrasts with other studies where overexpression of parkin led to a clear mitochondrial localization in other cell lines including neuronal cells [40,41,45,46]. Based on these findings, we analysed endogenous and EGFP-parkin translocation to mitochondria in two human immortalized neuroblastoma cell lines, BE(2)-M17 and SH-SY5Y. Similarly to differentiated striatal cells, endogenous parkin was not translocated to mitochondria in BE(2)-M17 or SH-SY5Y cell lines upon mild or acute depolarization (**Fig EV 1A**). In contrast, artificial overexpression of EFGP-parkin under strong depolarizing conditions led to nearly all cells showing EGFP-parkin translocation to mitochondria (**Fig EV 1B**). According to these results, and in agreement with other studies focused on neuronal mitophagy, endogenous parkin levels in striatal neurons seem too low for a clear parkin translocation to the mitochondria to be detected. Considering the latest model of PINK1/parkin-dependent mitophagy [30], parkin’s role in mitophagy is likely to be important as an amplifier of the mitophagy signal, but it is unlikely to be essential for mitophagy to take place. When parkin levels in mitochondria are low (below detectable levels), mitophagy can still take place through a PINK1-dependent process; however, the lack of parkin as an amplifier protein leads to a lower rate of polyubiquitination of the mitochondrial surface protein and consequently mitophagy occurs at a lower rate [30]. In our neuronal model, mitophagy probably takes place at a slower rate or in a less-robust manner, but still works since we were able to detect mitochondria engulfed by autophagosomes and fused with lysosomes (**Fig. 1A**). According to these results, we suggest that induced mitophagy taking place in striatal neurons is independent of parkin translocation to mitochondria, and that this event is not related to Htt protein. Indeed, the observed parkin-independent mitophagy seems to be related to the nature of the neuronal stratial cells themselves since we observed the same effect in both WT and mHtt cells. We can therefore postulate that parkin translocation does not occur in this cell type and mitophagy is taking place at a low level. The observed increase in basal PINK1 levels could be related to alternative roles of PINK1 in other mitochondrial quality control mechanisms that act as sensors of mitochondrial status [26,47,48] since mitochondrial dysfunction is present in HD as an early event of the disease and PINK1 not only detects mitochondria depolarization to trigger mitophagy, but also is involved in many other mitophagy-independent mechanisms that respond to mitochondrial damage.

**Figure 2.**
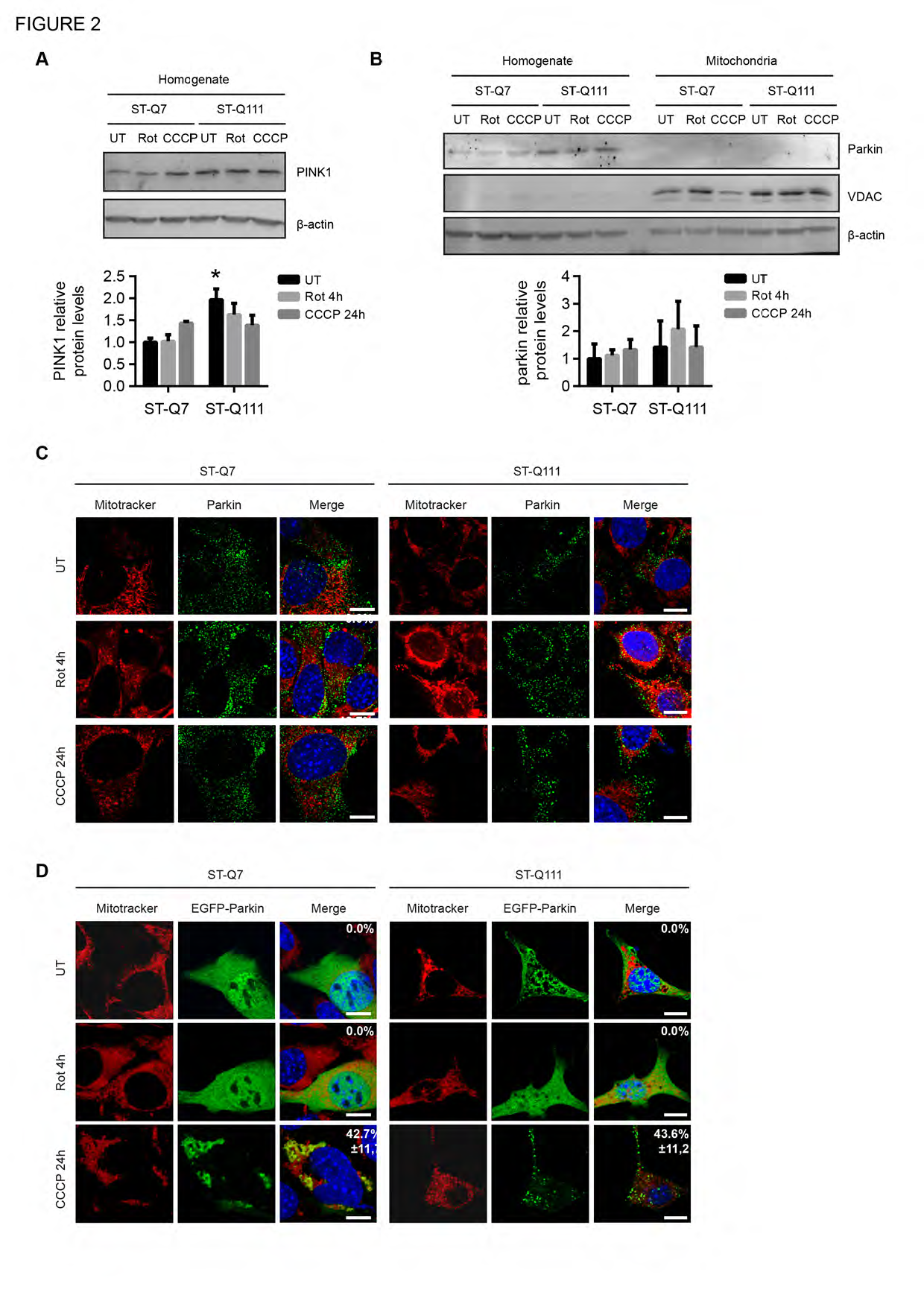
Mitophagy in differentiated striatal cells is not mediated by Parkin translocation to mitochondria. (**A**) Representative immunoblots of PINK1 protein levels in total homogenates from ST-Q7 and ST-Q111 cells untreated (UT) or treated with rotenone (Rot, 1 μM, 4h) or CCCP (10 μM, 24h). Protein levels were normalized relative to ß-actin and quantification is depicted as fold change to ST-Q7 untreated. **B**) Representative immunoblots of Parkin protein levels in total homogenates and isolated mitochondria from ST-Q7 and ST-Q111 cells untreated UT or treated with Rot (1 μM, 4h) or CCCP (10 μM, 24h). Protein levels were normalized relative to ß-actin and quantification is depicted as fold change to ST-Q7 untreated. (**C**) Representative images of MitoTracker^®^ DeepRed and Parkin immunostained ST-Q7 and ST-Q111 cells UT or treated with Rot (1 μM, 4h) or CCCP (10 μM, 24h). (**D**) Representative images of MitoTracker^®^ DeepRed and EGFP-Parkin immunostained ST-Q7 and ST-Q111 cells UT or treated with Rot (1 μM, 4h) or CCCP (10 μM, 24h). (**A-B**) Data are presented as mean ± s.e.m. from three independent experiments. ^*^ P < 0.05 compared to ST-Q7 untreated after two-way ANOVA test followed by Tukey’s post hoc test. **(C-D**) Scale bars: 10 μm. A minimum of cells 30 were analyzed per condition. Data are presented as mean ± s.e.m. of the percentage of cells with endogenous Parkin or EGFP-Parkin colocalizing with MitoTracker^®^ DeepRed labeled mitochondria.

### The polyQ tract in mutant Htt affects initation of the mitophagy process

Autophagy initiation is coordinated by ULK1, which forms a complex with Atg13 and FIP200 to regulate the initial step of autophagy induction in mammalian cells [49–51]. Under basal conditions, the ULK1 protein is bound to mTORC1 (mammalian target of rapamycin complex 1) and inactivated by mTOR-mediated phosphorylation at serine 757 (S757) [52]. To induce autophagy, ULK1 must be activated and released from mTORC1, losing the mTOR-mediated inhibiting phosphorylation at S757. Simultaneously, AMPK activates ULK1 through physical interaction wit it [53] and via the phosphorylation of the Ser555, Ser777, and Ser317 residues [52,54]. Previous studies showed that Htt plays a key role as a scaffold protein by binding active ULK1 at its C-terminal domain, by competing with mTORC1 to bind to ULK1, and promoting the initiation of autophagy [27,28]. To determine the ability of ULK1 to be released from mTORC1 and to understand how the presence of the polyQ tract with 111 glutamines might affect this step, we analyzed the interaction of ULK1 with mTOR and Htt. To this end, we used the Proximity Ligation Assay (PLA) technology since immunoprecipitation with the endogenous proteins was technically not feasible.

In differentiated ST-Q111 cells, the ULK1-mTOR interaction was more stable after depolarization compared to that seen in ST-Q7 cells, where the ULK1-mTOR interaction decreased upon depolarization to permit ULK1 activation (**Fig. 3A**). These results suggest that, in mutant cells, ULK1 remains more inactive and bound to mTOR than in WT cells. In a similar way, the ULK1-Htt interaction was increased in ST-Q7 cells after depolarization, but this interaction was not induced in ST-Q111 after depolarization, confirming that ULK1 has a reduced affinity for mHtt (**Fig. 3B**). In agreement with the PLA result, mTOR-mediated ULK1 phosphorylation at S757 was more abundant in a mHtt cell line and increased after depolarization, while AMPK-mediated phosphorylation at ULK1 Ser555 was increased after depolarization only in ST-Q7 cells (**Fig. EV 2**). We also observed by PLA that another activation site mediated by AMPK phosphorylation at ULK1 Ser317 is also increased in WT cells after depolarization but remained unchanged in mHtt cells (**Fig. 3C**). Taken together, these results suggest that the extended polyQ tract in mHtt is somehow interfering with the ULK1 shift from the mTORC1 complex to the Htt scaffolding complex, a necessary step to initiate autophagy and consequently mitophagy as well.

**Figure 3.**
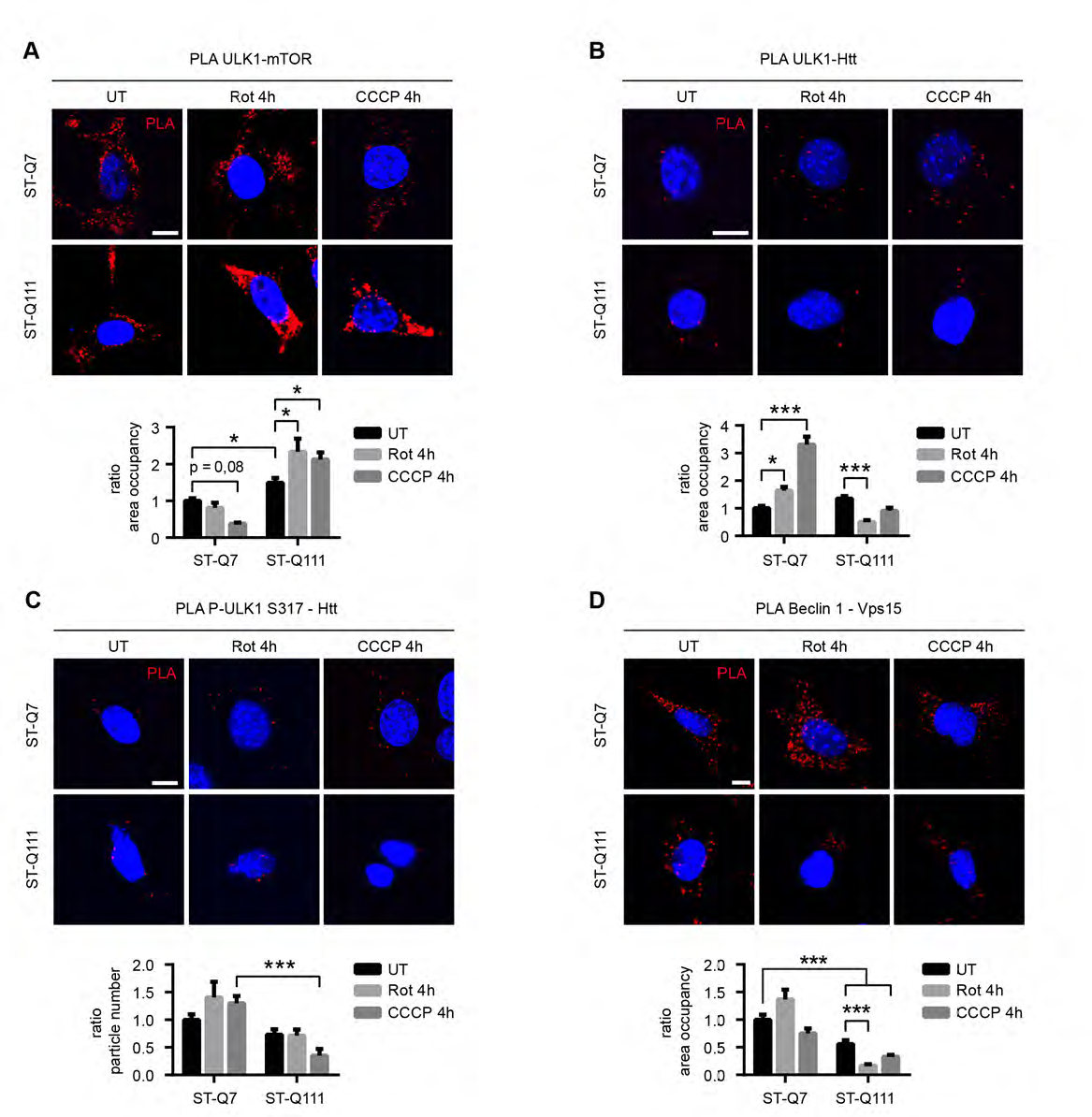
Mutant Htt alters the autophagy initiation step. (**A**) ULK1-mTOR interaction detected by PLA in ST-Q7 and ST-Q111 cells untreated (UT) or treated with rotenone (Rot, 1 μM, 4h) or CCCP (10 μM, 4h). 24 to 85 cells were analyzed per condition. (**B**) ULK1-Htt interaction detected by PLA in ST-Q7 and ST-Q111 cells UT or treated with Rot (1 μM, 4h) or CCCP (10 μM, 4h). (**C**) Phospho-ULK1 S317-Htt interaction detected by PLA in ST-Q7 and ST-Q111 cells UT or treated with Rot (1 μM, 4h) and CCCP (10 μM, 4h). **(D)** Beclin 1 - Vps15 interaction detected by PLA in ST-Q7 and ST-Q111 cells untreated (UT) or treated with Rot (1 μM, 4h) and CCCP (10 μM, 4h). 35 to 115 cells were analyzed per condition. Scale bars: 10 μm. PLA interaction was quantified as a ratio of area occupancy by the PLA signal or number of particles represented as the fold change compared with ST-Q7 UT condition. Data are presented as mean ± s.e.m. ^*^ P < 0.05, ^**^ P < 0.01, ^***^ P < 0.001 after two-way ANOVA test followed by Tukey’s post hoc test.

Following its activation, ULK1 phosphorylates beclin 1, a component of the autophagy initiation complex. The complex formed by beclin 1, Vps15, Vps34 and other proteins is responsible for autophagosome formation [55]. Recently it was shown that the deubiquitinase ataxin 3, which is altered in spinocerebellar ataxia type 3, interacts through its WT polyQ domain with beclin 1 and protects beclin 1 from proteasomal degradation. However, beclin 1 binds not only to ataxin 3 but to a greater extent to expanded polyQ tracts, such as the ones present on mHtt, thus competing with the physiological binding of ataxin 3 and promoting the degradation of beclin 1. Lower levels of beclin 1 are related to lower initiation complex assembly and a lower rate of macroautophagy [56]. Using PLA, we investigated whether this newly identified toxic function of mHtt could affect mitophagy at this step by impairing the formation of beclin 1-Vps15-Vps34 complex as one of the initial protein complexes required for the initiation of autophagy. At basal levels, the beclin 1-Vps15 interaction was downregulated in the presence of mHtt. Upon depolarization, differentiated ST-Q7 cells exhibited increased beclin 1-Vps15 interaction, indicative of the assembly of the beclin 1 initiation complex, while differentiated ST-Q111 cells displayed a downregulation of this complex formation (**Fig. 3D**). These results further confirm that expanded polyQ tracts present in mHtt impair formation of the autophagy initiation complex in striatal cells, a result that is in agreement with a previously described reduction of beclin 1 levels in different cellular and animal models with expanded polyQ forms [56]. In summary, mHtt affects the formation and stability of two complexes that are essential for autophagosome formation during autophagy initiation. This scenario might affect all forms of autophagy, but certainly the initiation of mitophagy will also be impaired due to the reduced assembly of these complexes.

### The polyQ tract in mHtt affects the scaffolding role of Htt during mitophagy

Previously, we and others have shown that, in addition to its interaction with ULK1, Htt can also interact with the the selective autophagy adaptor protein p62 [27,28,23]. Besides p62, different protein receptors for selective mitophagy like NDP52 (also known as CALCOCO2), optineurin (OPTN), NBR1 and TAX1BP1 have been identified [57]. During mitophagy, these receptors are able to recognize polyubiquitinated OMM proteins and, simultaneously, LC3-II in the autophagosome membrane. Whereas initially some studies showed the requirement of p62 for mitophagy [58,59], others have reported that lack of p62 does not impair mitophagy [60,61]. In a recent study using the combined knock-out of all five mitophagy receptors, it has been shown, that although all of the receptors can intervene during mitophagy, only two of them, OPTN and NDP52, are essentials [30] with the others probably having redundant roles. We focused on these two essential receptors with a view to determining whether they are also able to interact directly with the Htt scaffolding complex as previously observed with p62 [27,28,23] and NBR1 (data not shown) With our objective being to determine if this interaction could be affected by the polyQ tract, we first determined the presence of OPTN-Htt and NDP52-Htt interactions using PLA (**Fig. 4A-B**). Our results were in agreement with previous studies [62,63] and to protein-protein interaction predictions. Further to this, we detected a reduced OPTN-Htt interaction after depolarization, but only when Htt was carrying the mutant polyQ tract (**Fig. 4A)**; this is is in agreement with the decrease in OPTN-Htt colocalization reported in a previous study [62]. Moreover, NDP52-Htt interaction in the presence of mutant polyQ could not be detected (**Fig. 4B**). This absence of interaction between NPD52 and mHtt was not due to the absence of NDP52 protein but to the total loss of interaction, since NDP52 protein could still be detected by immunofluorescence in mHtt cells before and after depolarization (**Fig EV 3**). These results show that the interaction between the two essential receptors of mitophagy and Htt is influenced by the number of glutamines in the polyQ tract. In the case of NDP52, the interaction is totally abolished when mHtt is present.

**Figure 4.**
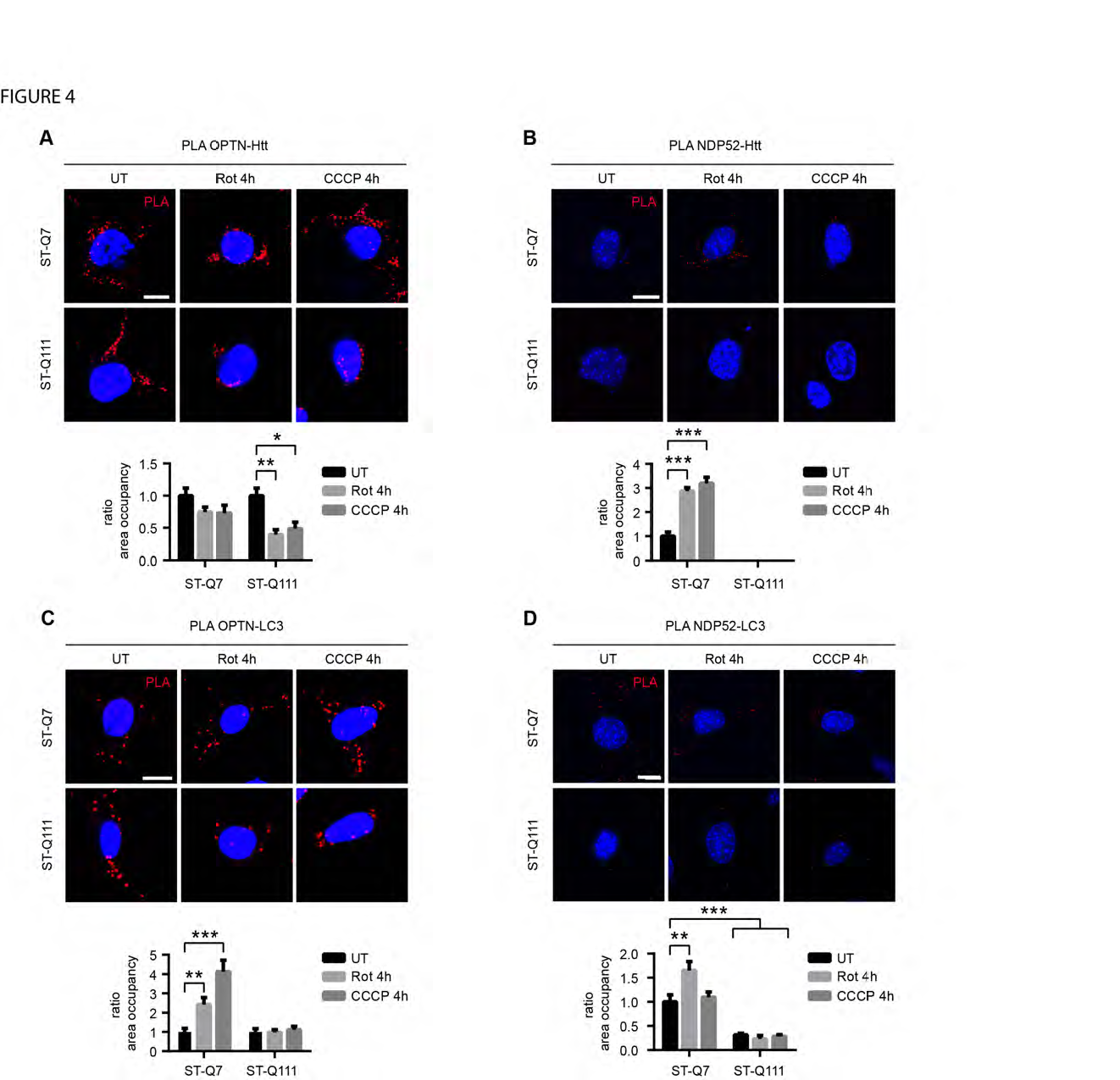
polyQ tract in mutant Htt affects the interactions with the mitophagy receptors and the recruitment of LC3-II. (**A**) OPTN-Htt interaction detected by PLA in ST-Q7 and ST-Q111 cells untreated (UT) or treated with rotenone (Rot, 1 μM, 4h) or CCCP (10 μM, 4h). 20 to 30 cells were analyzed per condition. (**B**) NDP52-Htt interaction detected by PLA in ST-Q7 and ST-Q111 cells UT or treated with Rot (1 μM, 4h) or CCCP (10 μM, 4h). 38 to 46 cells were analyzed per condition. (**C**) Representative images of NDP52 immunostained ST-Q7 and ST-Q111 cells UT or treated with CCCP (10 μM, 4h). (**C**) OPTN-LC3 interaction detected by PLA in ST-Q7 and ST-Q111 cells untreated (UT) or treated with rotenone (Rot, 1 μM, 4h) or CCCP (10 μM, 4h). 25 to 38 cells were analyzed per condition. (**D**) NDP52-LC3 interaction detected by PLA in ST-Q7 and ST-Q111 cells UT or treated with Rot (1 μM, 4h) or CCCP (10 μM, 4h). 25 to 45 cells were analyzed per condition. Scale bars: 10 μm. PLA interaction was quantified as a ratio of area occupancy by the PLA signal represented as the fold change compared with ST-Q7 untreated condition. Data are presented as mean ± s.e.m. ^*^ P < 0.05, ^**^ P < 0.01, ^***^ P < 0.001 after two-way ANOVA test followed by Tukey’s post hoc test.

As the last step of substrate recognition during mitophagy, receptors interact directly with and recruit LC3-II, thereby allowing elongation of the autophagosome membrane around mitochondria. We used PLA to analyze how the presence of the polyQ expansion could indirectly affect the binding/recruitment of LC3-II by the receptors to the mitophagy initiation site. We analyzed the interaction between the two essential mitophagy receptors (OPTN and NDP52) with LC3 and observed as expected that upon depolarization, the OPTN-LC3 interaction and the NDP52-LC3 interaction were increased in differentiated ST-Q7 cells. Conversely, in the presence of the mutant polyQ tract, the OPTN-LC3 interaction was impaired and the NDP52-LC3 interaction decreased, probably as a consequence of the failure of receptors to be recruited to the mitophagy site (**Fig. 4C and D**). In addition to the essential receptors, we also examined the interaction between the autophagy receptors p62 and NBRI with LC3-II by immunoprecipitation and observed the same effect; that mHtt dramatically impairs p62 and NBRI interaction with LC3-II (**Fig EV 4**). Taken together, these results confirm the pivotal role of Htt as a scaffold for several mitophagy mechanisms, including the last step of LC3-II recruitment by receptors to the mitophagy initiation site, a key step to promoting enlongation of the autophagosome membrane to engulf tagged mitochondria.

### Impaired mitophagy leads to accumulation of damaged mitochondria in ST-Q111 cells

As a consequence of decreased mitophagy efficiency after depolarization, an accumulation of damaged mitochondria can be observed in differentiated ST-Q111 neurons. The ratio between non-depolarized mitochondria, detected with a mitochondrial probe that is sensitive to the mitochondrial membrane potential (TMRM), and total mitochondria, detected with a mitochondrial membrane potential-insensitive mitochondrial probe (MitoTracker^®^ Deep Red) indicates the number of healthy mitochondria remaining after depolarization. We observed that this ratio was higher in control cells (ST-Q7), whereas mutant cells (ST-Q111) presented lower rates of healthy mitochondria after 24h of mild or acute depolarization (**Fig. 5A**). This finding indicates that mutant cells have a reduced capacity for turnover of damaged mitochondria. In other experiments, we used Seahorse XF technology to analyze mitochondrial function by measuring the oxygen consumption rate (OCR) of differentiated striatal cells under basal conditions and in response to cellular respiration stress. Initially, both differentiated neuronal cell lines had similar basal respiration levels (**Fig. 5B and C**).). Inhibition of ATP synthase (complex V) with oligomycin caused an expected decrease in respiration in both ST-Q7 and ST-Q111 cells with no significant difference between the two cell types. Addition of the uncoupling agent CCCP collapsed the proton gradient and caused an increase in the OCR until the maximum respiration rate was achieved. ST-Q111 cells clearly showed a lower rate of maximal respiration and a lower spare respiratory capacity compared with control cells (**Fig. 5B and C**), indicating that mitochondria from ST-Q111 cells are bioenergeticaly less active and have a decreased ability to respond to an increased energy demand. When cells were mildly depolarized by exposure to rotenone for 4h and then left to recover for 24h, the remaining mitochondria from mutant ST-Q111 cells exhibited lower levels of basal respiration, ATP production, maximal respiration and spare respiratory capacity compared to control cells subjected to the same treatment (**Fig. 5D**). This assay clearly shows that, owing to an inefficient induced mitophagy, the remaining pool of mitochondria present in the cell is not fully healthy, thus contributing to the general mitochondrial dysfunction observed in HD. One previous study also reported the presence of decreased spare respiratory capacity in mutant huntingtin striatal neurons [64] while others observed respiratory deficits only when respiration demand was high [65,66]. Interestingly, dysfunctional respiration was only observed in striatal neurons but not in cortical neurons [65] which degenerate later in the course of the disease. In the same way, several studies have also described dysregulated glycolysis in the context of HD [67–69]. One of the main consequences of the cell’s inability to properly eliminate damaged mitochondria is the increase in reactive oxygen species (ROS), produced at least in part from the damaged mitochondria. These can be detected by the fluorescent probes CellROX^®^ Green (**Fig. 5E and F**) and CM-H2DCFDA (**Fig 5G**). Our data showed increased ROS levels in ST-Q111 cells compared with ST-Q7 cells, which might be due in part to the decreased induced-mitophagy rate observed in mutant cells. Higher basal levels of mitochondria-generated ROS and mtDNA lesions were also reported in mutant huntingtin striatal cells [64], which correlated with the increased presence of oxidative damage in proteins and lipids in the striatum and cortex of human HD brains [70–73].

**Figure 5.**
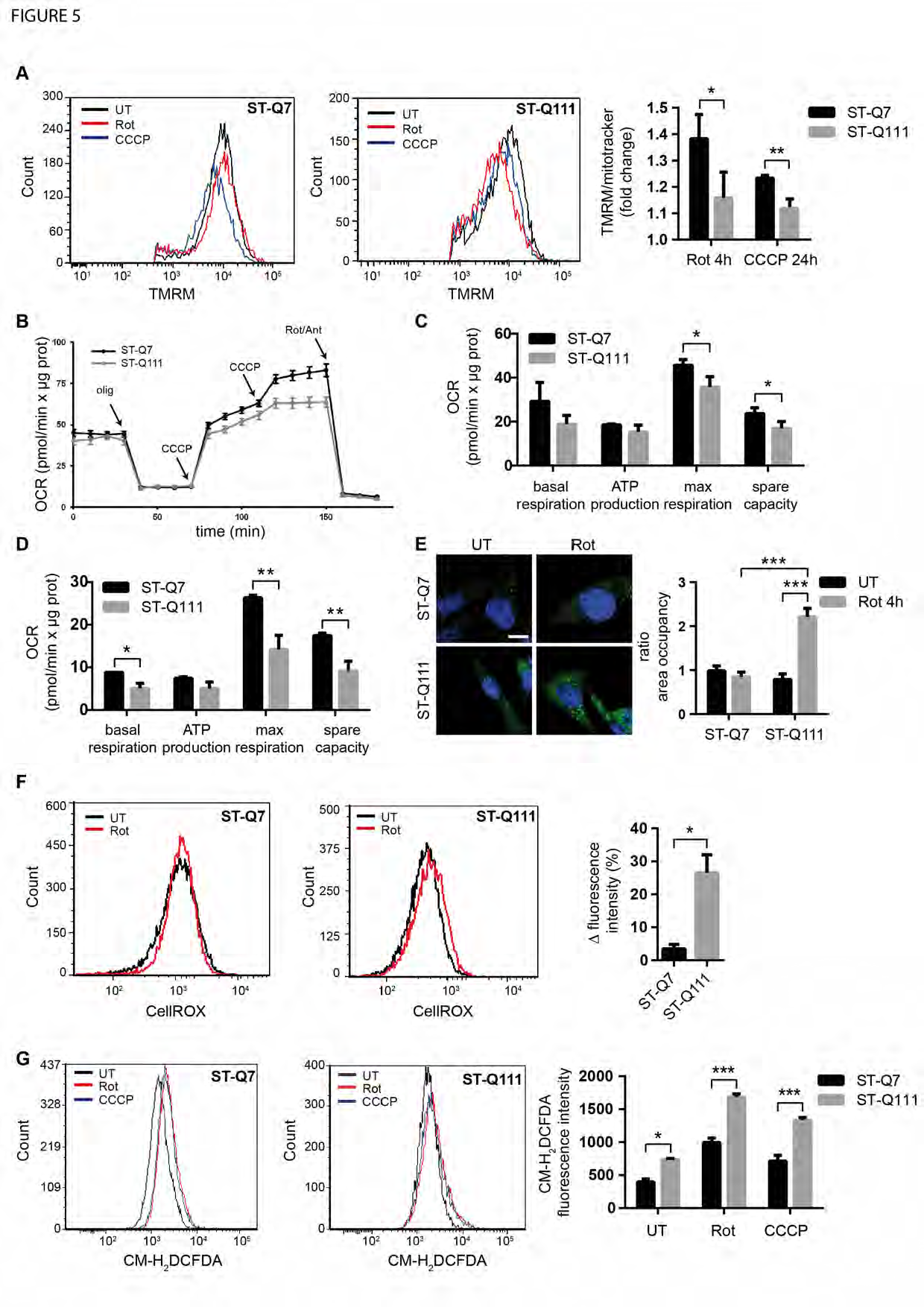
Impaired mitophagy leads to the accumulation of damaged mitochondria in ST-Q111 cells. (**A**) Flow cytometry was used to determine TMRM and MitoTracker^®^ DeepRed staining in ST-Q7 and ST-Q111 cells untreated (UT) or treated with rotenone (Rot, 1 μM, 4h) or CCCP (10 μM, 4h). Cells were added chloroquine with the rotenone and CCCP treatments (60 μM; n=3 independent experiments). Quantification is represented as fold change compared with the corresponding UT condition. (**B**) Seahorse XF-24 instrument analysis of oxygen consumption rate (OCR). Mitochondrial respiration reflected by OCR levels was detected in ST-Q7 and ST-Q111 cells under basal conditions or following the addition of oligomycin (olig, 1 μM), the uncoupler CCCP (CCCP, 0.5 μM each time) and Rotenone and Antimycin (Rot/Ant, 1 μM) at the indicated times (n=3 independent experiments). (**C**) The rates of basal respiration, ATP production, maximal respiration and spare capacity were quantified by normalization of OCR level to the total protein OD values (n=3 independent experiments). (**D**) The rates of basal respiration, ATP production, maximal respiration and spare capacity in ST-Q7 and ST-Q111 cells after 4h of rotenone treatment and 24h of recovery were quantified by normalization of OCR level to the total protein OD values. (**E**) Representative images of CellROX^®^ Green immunostained ST-Q7 and ST-Q111 cells UT or treated with Rot (1 μM, 4h). Scale bar: 10 μm. Intracellular ROS levels were quantified as a measure of the area fraction occupied by CellROX^®^ staining and represented as the fold change compared with ST-Q7 UT. 44 to 80 cells were analyzed per condition. (**F**) Flow cytometry was used to determine CellROX^®^ intensity in ST-Q7 and ST-Q111 cells UT or treated with Rot (1 μM, 4h). Intracellular ROS levels were quantified as a measure of CellROX^®^ intensity and represented as the ratio between rotenone and untreated conditions (n=3 independent experiments). (**G**) Flow cytometry was used to determine CM-H_2_DCFDA staining in ST-Q7 and ST-Q111 cells UT or treated with Rot (1 μM, 4h). Intracellular ROS levels were quantified as a measure of CM-H_2_DCFDA intensity and represented as mean fluorescence intensity (n=3 independent experiments). (**A-G**) Data are presented as mean ± s.e.m. ^*^ P < 0.05, ^**^ P < 0.01, ^***^ P < 0.001 after (A; E; G) two-way ANOVA test followed by Tukey’s post hoc test (C-D; F) Student’s t test.

In summary, this work has analysed in-depth the steps by which huntingtin protein modulates mitophagy under conditions that are as close as possible to physiological, with a view to identifying the consequences of the polyQ tract expansion. To this end, we have used differentiated striatal cells expressing endogenous levels of mitophagy-related proteins. In addition, we have used CCCP to induce a fast and massive mitochondrial depolarization, as well as rotenone at a low concentration, which induces a milder depolarization closer to that which may occur during the evolution of the disease. The results obtained in this work argue in favour of the presence of loss-of-function mechanisms in the HD pathology. By promoting the activation of mitophagy and the recruitment of different elements participating in this mechanism, our studies have demonstrated the key role of Htt protein in mitophagy. The presence of the mutation in Htt affects the ability of the protein to execute its role and consequently impairs the efficiency of the mitophagy process, leading to the accumulation of damaged mitochondria and the generation of ROS.

At the present we cannot rule out the possibility that other forms of mitophagy that have been described for different stress conditions and that are independent of ubiquitin labeling, PINK1/parkin function and cytosolic receptors [26,48] are not also taking place in these striatal cells after depolarization; these forms could be partially compensating for the effects of mHtt. While the possible role of Htt protein in these other, potentially redundant pathways is unknown and requires further research, the present work nevertheless demonstrates that Htt plays an important role in induction-mediated mitophagy during the initiation steps and in recruitment of the essential mitophagy receptors.

In addition of these effects on mitophagy, numerous other alterations to mitochondria homeostasis have also been reported in HD models and tissue, including bioenergetic alterations [74,75], reduced activity of OXPHOS complex I, II and IV [76–78], reduced ATP levels [79,80], increase of oxidative stress [81,73,72,82,83], and defects in mitochondrial dynamics [84–86], mitochondrial biogenesis [87–89], and mitochondrial trafficking [15,16,86,90,92–94].

Obviously not all these mitochondria-associated alterations are are likely to be due to Htt’s role in mitophagy,but they could occur as a consequence of other alternative Htt functions [95]. As the events occurring in HD neurons would likely accumulate over the time course of the development of the disease, and since mitophagy is one of the main mitochondrial quality control mechanisms that guarantees a healthy mitochondrial pool, the impairment of neuronal mitophagy described here must play an important role in the pathogenesis of the disease and contribute to the process of neurodegeneration.

## MATERIAL & METHODS

### Reagents

rotenone (Ref. R-8875), CCCP (carbonyl cyanide 3-chlorophenyl hydrazone, Ref. C2759), oligomycin (Ref. 75351), antimycin A (Ref. A8674) and the lysosomal inhibitors chloroquine (Ref. C6628) and bafilomycin A1 (Ref. B1793) were from Sigma-Aldrich. The following reagents were also used: MitoTracker™ Deep Red FM (Molecular Probes, Ref. M22426), CM-H_2_DCFDA (Life Technologies, Ref. C6827), Cell Rox^®^ Green Reagent (Thermo Fisher, Ref. C10444) and TMRM (Tetramethylrhodamine methyl ester, Sigma-Aldrich, Ref. T668).

### Antibodies

LC3 (NB-2220), PINK1 (BC100-494SS), ULK1 (NBP2-24738S) and Vps15 (NBP1-30463) from Novus, LAMP-1 (sc-19992), NDP52/CALCOCO2 (sc-376540) and Beclin 1 (sc-48341) from Santa Cruz, Parkin (ab15954), Parkin (ab77924) and VDAC1/Porin (ab15895) from Abcam, poly-UB K63 (5621S), ULK1 Ser555 (5869P), ULK1 Ser317 (12753S) and ULK1 Ser757 (6888S) from Cell Signaling, OPTN (10837-1-AP) from Proteintech, Htt (MAB2166) from Chemicon, p62 (GP62-C) and NBR1 (H00004077-M01) from Abnova, anti-DNA (AC-30-10) from Progen, mTOR [GT649] mouse (GTX630198) from Genetex, total-UB (U5379) and β-actin (A5441) from Sigma-Aldrich. Secondary antibodies goat anti-mouse HRP, donkey anti-rabbit HRP, donkey anti-rat HRP from GE Healthcare, goat anti-guinea pig HRP from Santa Cruz, Alexa Fluor^®^ 488 goat anti-mouse, Alexa Fluor^®^ 488 goat anti-rabbit, Alexa Fluor^®^ 488 donkey anti-rat, Alexa Fluor^®^ 594 goat anti-mouse and Alexa Fluor^®^ 594 goat anti-rabbit from Thermo Fisher.

### Cell culture and treatments

#### ST-Q7 and ST-Q111 cell lines

ST*Hdh*^Q7/Q7^ (ST-Q7) and ST*Hdh*^Q111/Q111^ (ST-Q111) were obtained from Coriell Institute (Ref. CH00097 and CH00095, respectively). ST-Q7 and ST-Q111 cells were grown in Dulbecco’s Modified Eagle Medium (DMEM, Gibco, Thermo Fisher, Ref. 41966-029) supplemented with 10% inactivated fetal bovine serum, 1% penicillin-streptomycin and 0.4 mg/mL active geneticin and maintained at 33°C in humidified 95% air/5%CO_2_ incubator. For differentiation, cells were supplemented with dopamine (DOPA) cocktail consisting of 10 ng/mL α-FGF-acidic mouse recombinant (Fibroblast growth factor, Sigma-Aldrich, Ref. SRP3197-50U), 250 μM IBMx (3-isobutyl-1-methylxanthine, Sigma-Aldrich, Ref. I5879), 200 nM TPA/PMA (Phorbol 12-myristate 13-acetate, Sigma-Aldrich, Ref. P1585), 50 μM Forskolin (Sigma Aldrich, Ref. F6886) and 20 μM dopamine hydrochloride (Sigma-Aldrich, Ref. H8502) during at least 24h.

#### BE(2)-M17

Human neuroblastoma dopaminergic cell line BE(2)-M17 (Sigma-Aldrich, Ref. 95011816) was grown in Minimal Essential Medium optimized (Opti-MEM, Gibco, Thermo Fisher, Ref. 31985062) supplemented with 10% inactivated fetal bovine serum (Thermo Fisher, Ref 26140-079), 1% penicillin-streptomycin (Sigma-Aldrich, Ref P4333) and 0.5 mg/mL active geneticin (Thermo Fisher, Ref 11558616) and maintained at 37°C in humidified 95% air/5%CO_2_ incubator.

#### SH-SY5Y

Human neuroblastoma SH-SY5Y cell line was obtained from ATCC (CRL-2266™) and grown in Dulbecco’s Modified Eagle Medium supplemented with 10% inactivated fetal bovine serum and 1% penicillin-streptomycin and maintained at 37°C in humidified 95% air/5%CO_2_ incubator.

#### Plasmid transfection

For plasmid transfection, Lipofectamine 3000 (Thermo Fisher, Ref. L3000015) was used. Cells were seeded in a 12-well plate and transfected with 0.8 μg (EGFP-Parkin) or 30 μg (mitoKeima) of DNA following the manufactures’s instructions. 6 hours after transfection, medium was changed to the corresponding growing medium.

#### Immunofluorescence microscopy

Cells were seeded in 16 mm of diameter coverslips (Menzel Gläser, Ref. #1). Cells were fixed in 4% paraformaldehyde for 30 minutes at RT. Following the incubation with blocking solution containing normal goat serum 3%, triton x-100, 0.1% in PBS for 1h at RT, the corresponding primary antibodies were used diluted in the same blocking solution overnight at 4°C. Then, cells were incubated 1h at RT with the corresponding Alexa secondary antibodies, diluted in blocking solution. Nuclei were stained with 10 μM Hoechst 33342 in PBS for 10 min at RT. When indicated, before fixing, 100 nM MitoTracker™ Deep Red FM prepared in Opti-MEM medium was added and cell incubated during 30 minutes and washed twice with PBS before proceeding with the protocol. Slides were mounted onto superfrost ultra plus slides using DAKO Fluorescent Mounting Medium. Images were acquired using an Olympus FV1000 confocal microscope. Intensity analysis was performed by drawing a ROI to each cell and measuring its intensity using ImageJ 1.50a. Colocalization analysis of LAMP1-mitochondria was done with single cells quantifying the overlapping signal of LAMP-1 and MitoTracker™ Deep Red FM in the cytosol area. For such, we used the Pearson’s correlation coefficient and the percentage of lysosomes that were positive for mitochondria by using ImageJ 1.50a software. At least 50 single cells were quantified in each condition.

#### Immunoblot detection

For biochemical studies cells were washed twice with ice-cold PBS solution containing 137 mM NaCl, 2.7 mM KCl, 10 mM Na_2_HPO_4_ and 1.8 mM KH_2_PO_4_ at pH 7.4 (Sigma-Aldrich, Ref. P9333, S7907 and P5655, respectively) before detaching them with a cellular scraper. Cell suspension was centrifuged at 800g for 5 min at 4°C. Supernatant was discarded and the pellet was resuspended in PBS. We repeated the centrifugation step two more times and finally cell suspension was resuspended in RIPA buffer to which we added freshly proteases inhibitors (HALT Protease inhibitor cocktail EDTA free, Thermo Scientific, Ref. 1862209) and 1 mM PMSF (Sigma Aldrich, Ref. P7626). Homogenates were incubated on ice for 15 min and samples were sonicated. Total protein concentration was determined by the bicinchoninic assay method with BSA as a standard protein. Cellular protein extracts homogenized were heated at 95°C for 5 min after adding loading buffer 6x containing Tris 500 mM, 30% glycerol (Sigma-Aldrich, Ref. G5516), 10% SDS and 0.6 M DTT (Roche, Ref. 10708984001). Proteins were resolved by SDS-PAGE on different percentage of polyacrylamide gels, ranging from 7% to 15% and run around 90 min at 120V in running buffer (25 mM Trizma R Base, 192 mM glycine and 1% SDS, Sigma-Aldrich, Ref. T6066, G7126 and L3771, respectively). Protein All Blue Standards ladder (Biorad, Ref. 161-0373) was used as a ladder. Resolved proteins were transferred onto nitrocellulose membranes (GE Healthcare, Ref. 10600002) during 90 minutes at 200 mA per gel in transfer buffer (25mM Trizma R Base, 192mM glycine and 20% methanol) and then blocked with 5% non-fat milk powder (Sigma-Aldrich, Ref. 70166) in PBS for 1h at room temperature. Membranes were incubated with the corresponding primary antibodies diluted in 4% bovine serum albumin (Sigma-Aldrich, Ref. A4503) in PBS overnight at 4°C. Then, we proceeded to the incubation with the corresponding secondary antibodies coupled with horseradish peroxidase and diluted in 5% milk in PBS for 1h at RT. Finally, proteins were visualized using either West Pico SuperSignal Substrate or SuperSignal West Femto (Thermo Fisher, Ref. 34080 and 34095, respectively) on an ImageQuant RT ECL imaging system (GE Healthcare). Immunoblots were quantified by densitometry using ImageJ 1.50a.

#### Mitochondrial enrichment

Mitochondria were isolated from BE(2)-M17 cells as previously described on Frezza *et al*. [96]. Briefly, cells were washed once with PBS and detached them using a cell scraper. Cell suspension was centrifuged at 600g at 4°C for 5 min and the supernatant was discarded. Pellet was resuspended in ice-cold IBc buffer (10 mM Tris-MOPS (Sigma, Ref. M5162), 1mM EGTA/Tris (Sigma, Ref. E4378) and 0.2 M sucrose in H_2_O at pH 7.4). To homogenize we used a Teflon pestle operated at 1600 rpm and stroked the cell suspension placed in a glass potter during 30-40 times. Cell suspension was centrifuged at 600g for 10 min at 4°C and the supernatant was further centrifuged at 7000g for 10 min at 4°C. Pellet was washed with ice-cold IBc buffer and centrifuged again at 7000g for 10 min at 4°C. Finally, the pellet containing mitochondria was resuspended in an adequate volume of IBc buffer. Next, we determined the mitochondrial protein concentration by the bicinchoninic assay method with BSA as a standard protein.

#### Mitochondrial membrane potential

ST-Q7 and ST-Q111 cells were seeded and differentiated in 12-well plates and, after indicated treatments, cells were washed twice with pre-warmed PBS and 50 nMTMRM or 10 nM MitoTracker™ Deep Red FM probes were added to the medium and incubated for 30 min in cell incubator. Cells were washed twice with pre-warmed PBS and recollected with a cell scraper in medium. Using a FACSAria flow cytometer (BD Biosciences) 10.000 cells were acquired and fluorescence detected with the indicated lasers and detectors. Membrane potential was analyzed as mean fluorescence intensity (MFI) and represented either as MFI or as fold change compared to the corresponding controls using the FCS Express (v3 De Novo Software™).

#### Seahorse XF24 mitochondrial respiration analysis

Oxygen consumption rate (OCR) was measured using the Seahorse XF24 equipment (Seahorse bioscience Inc., North Billerica, MA, USA). ST-Q7 and ST-Q111 cells were seeded in Seahorse XF24 Cell Culture Microplate (Seahorse Biosciences, Ref. 100777-004). Striatal cells were seeded on XF24 microplates and differentiated and treated with the corresponding treatments. Cells were rinsed once and incubated in 700 μl of XF assay buffer (DMEM without NaHCO_3_, 2 mM glutamax; pH 7.4, 5 mM glucose, 1 mM sodium pyruvate), then equilibrated for 1 h at 37°C in a non-CO_2_ incubator. All medium and solutions of mitochondrial complex inhibitors were adjusted to pH 7.4 on the day of assay. Following three baseline measurements of OCR, mitochondrial complex inhibitors were sequentially injected into each well. Three OCR readings were taken after addition of each inhibitor and before automated injection of the subsequent inhibitor. Mitochondrial complex inhibitors, in order of injection, included oligomycin (1 μM) to inhibit complex V (i.e., ATP synthase), CCCP (0.5 μM each time) to uncouple the proton gradient, antimycin A (1.0 μM) to inhibit complex III, and rotenone (1.0 μM) to inhibit complex I. OCR were automatically calculated, recorded, and plotted by Seahorse XF24 software version 1.8 (Seahorse Bioscience, Billerica, MA, USA). At the end of each assay, cells were washed once with an excess of RT DPBS, lysed with ice-cold RIPA buffer and the protein content estimated by the bicinchoninic assay method with BSA as a standard protein.

#### Quantification of reactive oxygen species (ROS) by fluorescent probes

cells were seeded in 12-well plates and, upon 80% confluence, treated with the corresponding reagents. Cells were washed twice with pre-warmed PBS and either 50 μM of 2 μMCM-H_2_DCFDA probe or 50 μM CellROX was added in medium and incubated for 30 min in a cell incubator. For flow cytometry analysis, cells were washed twice with pre-warmed PBS and recollected with a cell scraper in medium. Using a FACSAria flow cytometer, 10.000 cells were acquired and fluorescence was read at 488 nm laser with 530/30 emission filter. ROS was analyzed as mean fluorescence intensity (MFI) and represented either as MFI or as a percentage compared to the corresponding controls using the FCS Express v3. For immunofluorescence analysis, cells were seeded in 16 mm of diameter coverslips and 5 μM of CellROX was added in medium and incubated for 30 min in a cell incubator. We followed the same protocol as the one for immunofluorescence. Images were acquired using an Olympus FV1000 confocal microscope.

#### Proximity Ligation Assat (PLA)

PLA was performed in 3% formaldehyde-fixed cells (Sigma Aldrich, Ref. F1635) following manufacturer’s instructions. Briefly, samples were blocked for 30 min at RT and incubated with specific primary antibodies overnight. Secondary antibodies anti-mouse PLUS and anti-rabbit MINUS conjugated with oligonucleotides were added to the reaction (Duolink PLA Probe anti-mouse PLUS and antirabbit MINUS, Duolink, Sigma Aldrich, Ref. DUO92001 and DUO92005 respectively) and incubated for 1h at 37°C. Ligation reagents were added and incubated for 30 min at 37°C. If the two PLA probes are close enough, a closed oligonucleotide circle will be generated. Amplification solution, containing nucleotides and fluorescently labeled oligonucleotides, and polymerase were added and incubated for 100 min at 37°C. The oligonucleotide arm of one of the PLA probes acts as a primer for rolling-circle amplification reaction using the ligated circle as a template. This generates a concatemeric product where the fluorescently labeled oligonucleotides will hybridize (Detection Reagent RED, Duolink, Sigma Aldrich, Ref. DUO92008). The proximity ligation signal is observed as fluorescence spots and was analysed by confocal microscopy using an Olympus FV1000 confocal microscope. The corresponding control experiments were performed under identical experimental conditions. Quantification was performed by analyzing the number of dots per cell and the area fraction that these dots occupy per cell using ImageJ 1.50a.

#### Immunoprecipitation

ST-Q7 and ST-Q111 cells seeded in 6-well plates, upon 80% confluence, were recollected in PBS using a cell scraper and centrifuged at 800g at 4°C for 5 min. Pellet was resuspended in IP buffer containing 0.01% triton x-100 in PBS and protease inhibitors (Thermo Scientific, Ref. 78415). 1:2 mixture of Dynabeads^®^ protein A and protein G (Thermo Scientific, Ref. 10001D and 10003D, respectively) was incubated with the corresponding antibody, together with 5 mM BS^3^ (bis(sulfosuccinimidyl)suberate, Sigma Aldrich, Ref. S5799), to crosslink the antibody to the Dynabeads^®^, in IP buffer and let rocking 30 min at RT. The supernatant was discarded using a magnetic particle concentrator (DynaMag™-2, Invitrogen, Ref. 4211) and BS^3^ was inactivated with 50mM Tris-HCl pH 7.4 for 15 min. Dynabeads^®^ were washed three times with IP buffer and incubated with the corresponding cell lysate for 90 min at 4°C. For elution, samples were incubated in elution buffer containing 0.2 M glycine pH 2.5, LB6x and 16 mM DTT 10 min at 70°C. Immunoprecipitates or whole-cell extracts were analyzed by standard immunoblotting.

#### Statistical analysis

the values were expressed as the mean ± standard error of the mean (SEM). The significant differences (^*^p<0.05, ^**^p<0.01, ^***^p<0.001) when comparing two groups were determined by a two-tailed unpaired Student’s t test. When comparing more than two groups, significant differences were determined by one-way or two-way ANOVA followed by Tukey’s post hoc test. Statistical analyses were performed using GraphPad Prism 6.

## ACKNOWLEDGEMENTS

This work was supported by Fondo de Investigación Sanitaria-Instituto de Salud Carlos III (Spain)-FEDER (CP09/00184, PI14/01529, PI17/00496, MSII15/00007), MINECO SAF2009-08374, and CIBERNED. S.F.I. was supported by an FPU doctoral fellowship from MINECO (Spain). A.P.Z. was supported by a Juan de la Cierva post-doctoral fellowship from MINECO (Spain).

## AUTHOR CONTRIBUTIONS

S.F-I performed immunofluorescence microscopy, immunoblot detection, PLA and FACS analysis, contributed to the data analysis and manuscript preparation. A.P-Z assited in immunofluorescence microscopy. D.S assited in Seahorse XF24 mitochondrial respiration analysis. M.V. assited in manuscript discussion; M.M-V. conceived the study, designed the experiments, performed immunofluorescence microscopy, FACS analysis and immunoprecipitation, analyzed data and wrote the paper.

## Extented figures

**Figure EV-1: (A)** Representative images of MitoTracker^®^ DeepRed and endogenous Parkin immunostained SH-SY5Y and BE(2)-M17 cells UT or treated with Rot (1 μM, 4h) or CCCP (10 μM, 24h).

**(B)** Representative images of MitoTracker^®^ DeepRed and EGFP-Parkin immunostained SH-SY5Y and BE(2)-M17 cells UT or treated with Rot (1 μM, 4h) or CCCP (10 μM, 24h). Scale bars: 10 μm. A minimum of cells 30 were analyzed per condition. Data are presented as mean ± s.e.m. of the percentage of cells with endogenous Parkin or EGFP-Parkin colocalizing with MitoTracker^®^ DeepRed labeled mitochondria.

**Figure Figure EV-2:** Representative immunoblots of Phospho-ULK1-S757, Phospho-ULK1-S555, total ULK1 (T-ULK1) and β-actin protein levels in total homogenates from ST-Q7 and ST-Q111 cells UT or treated with Rot (1 μM, 4h) or CCCP (10 μM, 24h). (**D**) Phospho-ULK1 S317-Htt interaction detected by PLA in ST-Q7 and ST-Q111 cells UT or treated with Rot (1 μM, 4h) and CCCP (10 μM, 4h).

**Figure EV-3:** Representative images of NDP52 immunostained ST-Q7 and ST-Q111 cells UT or treated with CCCP (10 μM, 4h). Scale bars: 10 μm.

**Figure EV-4:** Representative images of immunoprecipitation of p62 and NBR1 in ST-Q7 and ST-Q111 and immunoblotting against LC3-II.

**Figure EV-5:** Schematic representation of Htt’s role during mitophagy initiation. (**A**) WT Htt promotes the physical proximity of different protein complexes during initiation of the mitophagy process: Htt is needed to promote ULK1 activation by competing with the inhibitory complex mTORC1; Htt also recruites OPTN and NDP52 to the mitophagy initiation site and facilitates their interaction with LC3-II, the main component of the autophagosome membrane. (**B**) When mitophagy is induced in the presence of mHttt several steps pf mitophagy initiation are affected (red dashed arrows), ULK1 binds preferentially to mTOR, avoiding the activation of the ULK1 complex. mHtt directly interacts with beclin 1 impairing the formation of the Beclin1-Vps34 complex necessary for the initiation of selective autophagy. The two essential mitophagy adapters (OPTN and NDP52) are not recruited by mHtt and their interaction with LC3-II is impaired. Overall mitochondria are less efficiently engulfed by the autophagosomes and degraded in the lysosomes leading to the accumulation of damaged mitochondria.

